# The first complete mitochondrial genome of *Diadema antillarum* (Diadematoida, Diadematidae)

**DOI:** 10.1101/2022.10.05.510842

**Authors:** Audrey J. Majeske, Alejandro J. Mercado Capote, Aleksey Komissarov, Anna Bogdanova, Nikolaos V. Schizas, Stephanie O. Castro Márquez, Kenneth Hilkert, Walter Wolfsberger, Tarás K. Oleksyk

**Affiliations:** University of Puerto Rico at Mayagüez, Department of Biology, Call Box 9000, Mayagüez, PR 00681, U.S.A.; Oakland University, Department of Biological Sciences, 118 Library Drive, Rochester, MI 48309, U.S.A.; Applied Genomics Laboratory, SCAMT Institute, ITMO University, Lomonosova 9 Str., Saint Petersburg 197101, Russia; University of Puerto Rico at Mayagüez, Department of Marine Sciences, Call Box 9000, Mayagüez, PR 00681, U.S.A.

## Abstract

The mitochondrial genome of the long-spined black sea urchin, *Diadema antillarum*, was sequenced using Illumina next generation sequencing technology. The complete mitogenome is 15,708 bp in length, containing 2 rRNA, 22 tRNA and 13 protein-coding genes, plus a non-coding control region of 133 bp. The nucleotide composition includes 18.37% G, 23.79% C, 26.84% A and 30.99% T. The A + T bias is 57.84%. Phylogenetic analysis based on 12 complete mitochondrial genomes of sea urchins including four species of the family Diadematidae supported familial monophyly, however the two *Diadema* species, *D. antillarum* and *D. setosum* were not recovered as sister taxa.

## 1. Introduction

The long-spined black sea urchin, *Diadema antillarum* (Diadematoida, Diadematidae), is a marine benthic invertebrate that inhabits the shallow waters of the Western Atlantic Ocean and Caribbean Sea. It is an important herbivore and keystone species that helps maintain healthy coral reef systems along its coastal marine habitats (Liddell and Ohlhorst, 1986; Hughes et al., 1987; Carpenter, 1988, 1990; Hughes, 1994; Ferrari Legorreta, 2012; Vega Thurber et al., 2012; Burkepile et al., 2013; Steneck, 2020). After the human impact of overharvesting larger herbivorous fishes and the disappearance of larger vertebrates, *D. antillarum* were plentiful. Along with other smaller herbivore fishes they served as the primary grazers maintaining the health of a coral dominated reef system, up until the mid-1980s (Jackson, 2001; Pandolfi et al., 2003; Steneck, 2020). After the historic 1983–84 die-off event of this species presumably due to an unknown water borne pathogen, repeated die-off events occurred in the 1990s as well as in the current year (February - May, 2022, https://www.agrra.org/sea-urchin-die-off/; accessed May 24, 2022). These events have collectively decimated, as well as wiped out populations in some localities across the Caribbean (Lessios, 1995; Lessios, 1988; Lessios, 2016). Monitoring and recovery efforts have been promising, but future die-off events will likely occur as global climate change continues (Hylkema et al., 2022; Pilnick et al., 2021; Williams, 2022).

Until now, only a few genes have been described in the mitochondrial genome of *D. antillarum*, including partial sequences of *COI, COII* and *ATP6*, as well as assembled sequences of *ATP8* and *tRNA*^*Lys*^ (Lessios et al., 2001; Collin et al., 2020). While over 40 mitochondrial genomes of different sea urchin species have been completed so far, this includes only three species within the family Diadematidae, which contain 12 genera. Of these, both species of the genus *Echinothrix* have complete mitochondrial genomes in addition to one of the seven species within the genus *Diadema*, specifically *D. setosum*. In this study, we report the complete mitochondrial genome of *D. antillarum* and we explore the utility of whole mitochondrial genomes to place *D. antillarum* phylogenetically in the Diadematidae tree.

## 2. Data description

### 2.1 Sample collection and DNA extraction

An adult sea urchin was collected in Puerto Rico (18°20’35.2”N 67°15’40.5”W). A sample containing whole coelomocytes was withdrawn from the animal through the Aristotle’s Lantern using a sterile 23-gauge needle connected to a 5 mL syringe. The animal was photographed and then returned to its habitat at the collection site, thus, it was not retained as a voucher specimen for this study. The sample was held on ice during transport to the University of Puerto Rico Mayagüez (UPRM). Total DNA was extracted from coelomocytes per company instructions using an a DNeasy^®^ Blood and Tissue Kit (Qiagen, Inc.). The sample quality and concentration were assessed using a NanoDrop 2000 spectrophotometer (Thermo Fisher Scientific) prior to shipment for sequencing at the Psomagen (Macrogen USA) laboratory in Rockville, Maryland. Approval for the sample collection was obtained from the Department of Natural and Environmental Resources of Puerto Rico (O-VS-PVS15-AG-00046-01082018).

### 2.2 Sequencing and bioinformatics analysis

To prepare the DNA sample for sequencing, a library of sequences was generated using a TruSeq^®^ DNA PCR Free (350) library preparation kit (Illumina, Inc.). Quality and quantitation verification of libraries was performed using a Bioanalyzer. Next, an Illumina HiSeq 2500 platform generated paired-end 151 bp sequence reads. The sequencer produced 60.8 GB of total reads and 402,874,618 paired-end reads of which 88.21% had a quality score of ≥ Q30.

The raw read candidates for the mitochondrial genome were extracted bioinformatically from the total Illumina data that represented both nuclear and mitochondrial genomes using Cookiecutter software (Starostina et al., 2015). The reads were trimmed using a custom program called v2trim. Next, the list of kmers was formed in two stages. In the first stage we used the best-known reference genome of sea urchins (*Strongylocentrotus purpuratus*, NC_001453.1, Jacobs et al., 1988) to assemble the draft mtDNA contigs of *D. antillarum* using SPAdes genome assembler. In stage two, the assembled contigs were searched with BLAST NCBI web interface to find the closest reference sequence, which matched to the species, *Echinothrix diadema*, with the accession number KX385836.1. In the second stage, we constructed the kmer list using this closest reference sequence. The extracted reads were then assembled with SPAdes v3.15.4 using the default parameters (Bankevich et al., 2012). To verify assembly quality the extracted reads were mapped back to the full length mtDNA assembly with bwa mem using the default parameters (Li, 2013). The coverage plot was computed with bedtools genomecov with -d and - split parameters (Aaron et al., 2010). In addition, a per-base coverage data table was generated using the NGS data analysis tools provided in the Unipro UGENE software program (Okonechnikov et al. 2012). A graph of this data table was generated in Microsoft Excel (2019) and is presented as **Supplemental Figure 1**. Finally, the assembly start was rotated to tRNA-Phe. Following assembly, an online annotation was performed using the MITOS web server (http://mitos.bioinf.uni-leipzig.de/index.py) with following manual verification of each predicted RNA and protein. The results from this annotation were used to generate a mitochondrial map in Geneious Prime v2022.1.1 (**Figure 1**). The extracted reads, bam,coverage files and the reproducible commands list are available in our GitHub repository: https://github.com/aglabx/mtDNA_assembly/tree/master/Diadema_anthilarum.

**Figure 1.**
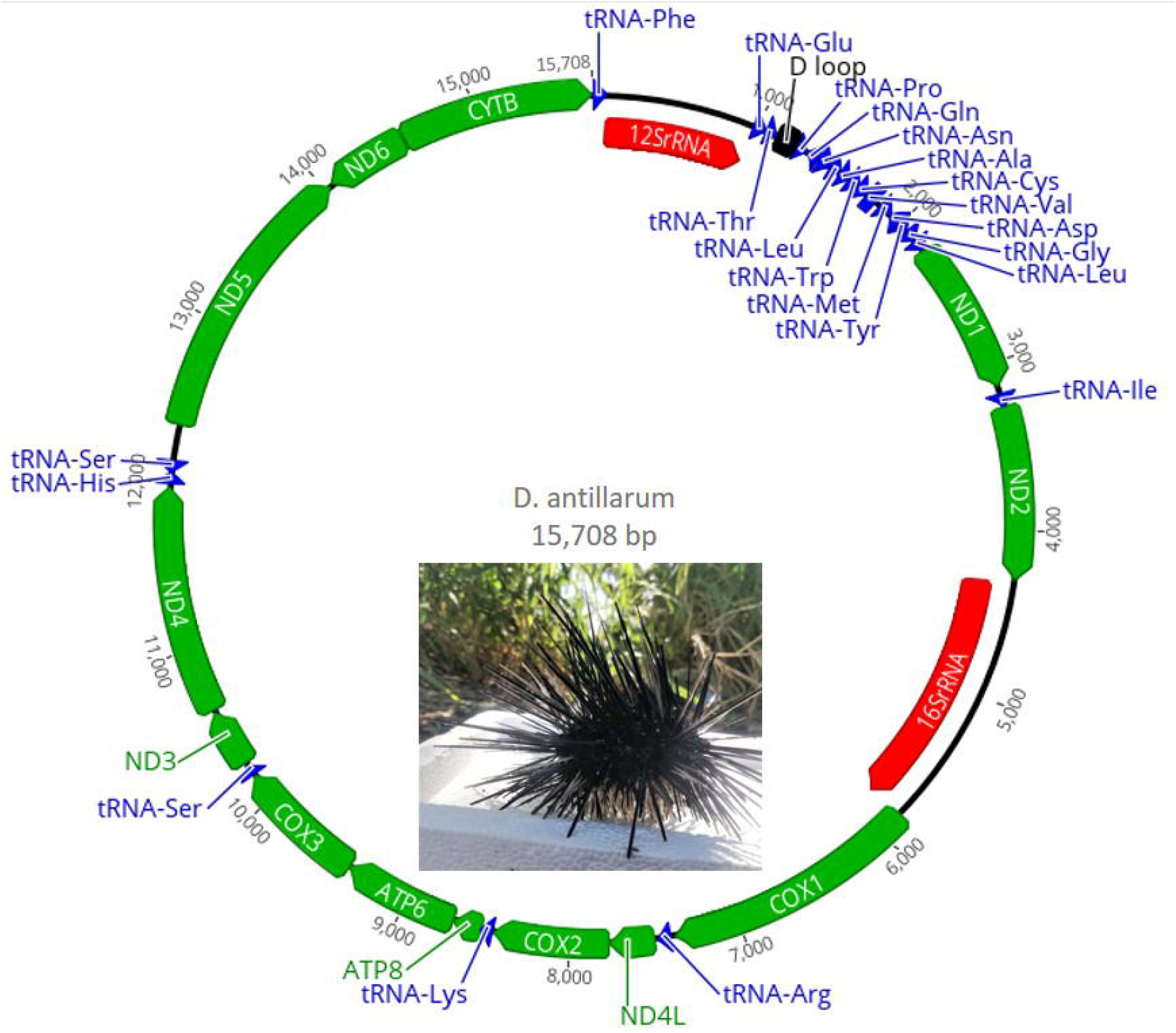
Mitochondrial genome map of *Diadema antillarum*. Arrow shapes are colored to indicate tRNA genes (blue), rRNA genes (red) and protein coding genes (green). The black rectangular shape refers to the non-coding control region or D loop. The directions of the arrows indicate direction of transcription on the H-strand (arrow points to the right) and L-strand (arrow points to the left). The map was generated in Geneious Prime v2022.1.1. The animal image is the specimen used for sampling in this study. Image taken by Alejandro Mercado Capote.

### 2.3 Phylogenetic analysis

Alignment and phylogenetic analysis were constructed in MEGA v11.0.11 (Tamura et al. 2021). Analysis included the complete mitochondrial genomes of 12 representative species from the orders Camarodonta (families Temnopleuridae, Echinometridae and Parechinidae), Arbacioida (family Arbaciidae), Temnopleuroida (family Toxopneustidae), Cidaroida (family Cidaridae), Echinoida (family Strongylocentrotidae), and Diadematoida (family Diadematidae) plus the sea cucumber, *Apostichopus* japonicus. A MUSCLE alignment method was implemented using the default parameters in the MEGA11 program. Another sequence alignment editor, BioEdit, was used to generate a sequence identity matrix of the aligned sequences (Hall, T.A., 1999). We performed an IQ-TREE analysis to determine the best-fit model of substitution among the mitogenome sequences (Nguyen et al 2015). A maximum likelihood tree (ML) was generated with the mitogenome nucleotide DNA sequences of the 12 sea urchin species and the sea cucumber that was included as an outgroup. We settled upon a comprehensive model of parameters for the phylogenetic analysis that resulted in branches containing the best supported bootstrap values (out of 500 bootstrap iterations) for the resulting consensus tree. The ML tree was generated using a Tamura-Nei model of evolution (Tamura and Nei, 1993). A discrete Gamma distribution was used to model evolutionary rate differences among sites (5 categories (+*G*, parameter = 0.3129)). Initial tree(s) for the heuristic search were obtained automatically by applying Neighbor-Join and BioNJ algorithms to a matrix of pairwise distances estimated using the Tamura-Nei model, and then the ML tree was generated by selecting the topology with superior log likelihood value. Additional sequence alignments and phylogenetic analysis was performed on available sequences for *Echinothrix* spp. and *Diadema* spp. for each of the three genes: 16S ribosomal RNA (partial sequence), ATPase genes (including ATP synthase subunit 6 gene partial coding sequence and tRNA-Lys gene partial sequence ATP8 gene complete coding sequence and ATP6 gene partial coding sequence) and cytochrome oxidase subunit 1 (CO1) (partial coding sequence). For these additional analyses, a MUSCLE alignment was generated using the default parameters in the MEGA11 program. In addition, we extracted the portions of the mitogenome for *E. diadema* (KX385836), *D. setosum* (KX385835) and *D. antillarum* (present study) that corresponded to each gene, by generating an initial alignment using available sequences from the same genera for each of the three genes. A ML tree was obtained for each gene using a bootstrapped (500 iterations) general time reversible (GTR) model with invariant (I) substitution rates among the nucleotides. The initial trees were obtained by applying Neighbor-Join and BioNJ algorithms to a matrix of pairwise distances estimated using the Tamura-Nei model, and then the ML tree was generated by selecting the topology with superior log likelihood value.

## Results

### 3.1 Signatures of the mitogenome

The circular mitogenome of *D. antillarum* is 15,708 bp in length, which consists of 2 rRNA genes, 22 tRNA genes, a non-coding control region, and 13 protein-coding genes that are common in other echinoderms, as well as the order of the genes (Bronstein and Kroh 1999, Ketchum et. al. 2018, De Giorgi et. al. 1996). Most of the genes are encoded on the H-strand except for one protein-coding gene, *ND6*, and five tRNA genes, including *tRNAGln, tRNAAla, tRNAVal, tRNAAsp*, and *tRNASer*, which are encoded on the L-strand (Bernt et al. 2013). Most of the protein-coding genes use the start codon ATG, with the exception of the *ATP8* gene, which starts with GTG. The length of the rRNA genes is 896 bp for *12S rRNA* and 1555 bp *16S rRNA*. The control region is 133 bp in length and contains the typical G repeat that is found in other echinoderms. This non-coding region is located at base positions 1111 to 1243, and is positioned in between the genes *tRNA*^*Thr*^ and *tRNA*^*Pro*^.

The composition of the nucleotides for the mitogenome of *D. antillarum* was calculated in the MEGA11 program, and includes 18.37% G, 23.79% C, 26.84% A and 30.99% T. The A + T bias was also calculated in MEGA, which is 57.84% for *D. antillarum*, slightly lower but comparable to that of *D. setosum*, which is 58.19%. In comparison to the two other *Echinothrix* species in the same family as *D. antillarum*, the A + T bias in the mitogenome of *D. antillarum* is slightly higher than that of *E. diadema* (57.61%) and *E. calamaris* (56.43%). Nonetheless, when compared to species in different orders, the mitochondrial A + T bias of *D. antillarum* is generally lower than other species in the following orders: Cidaroida (*Stylocidaris reini*, 59.93%; *Prionocidaris baculosa*, 59.14%; *Eucidaris tribuloides*, 59.70%), Camarodonta (*Echinometra mathaei*, 59.21%; *Heterocentrotus mamillatus*, 58.91%; *Heliocidaris crassispina*, 58.89%), and Echinoida (*Strongylocentrotus droebachiensis*, 58.96%; *S. intermedius*, 58.92%; *S. purpuratus*, 58.98%).

### 3.2 Phylogenetic tree

The bootstrap consensus ML tree is shown in **Figure 2**, which was inferred from 500 replicates through clustering of the associated taxa (Felsenstein, 1985). The results of the IQ-TREE analysis indicated that the best model of substitution was the most comprehensive model tested, the GTR+F+R2 model (General time reversible model with unequal rates and unequal base frequency with FreeRate rate heterogeneity model) (Tavaŕe 1986). The results of this test included a ML tree lacking a bootstrap analysis, which is available on the GitHub repository site listed above. In addition, a ML tree was generated in the MEGA11 program using the suggested GTR model of evolution with invariant (or unequal) substitution rates among sites (tree not shown). The two resulting ML consensus trees from the MEGA11 program plus the ML tree generated by the IQ-TREE analysis had identical topologies with regards to the species within the family Diadematidae. Of the two bootstrapped ML trees produced in MEGA, the tree presented in this study generally showed higher bootstrap values for all branches within the tree.

**Figure 2.**
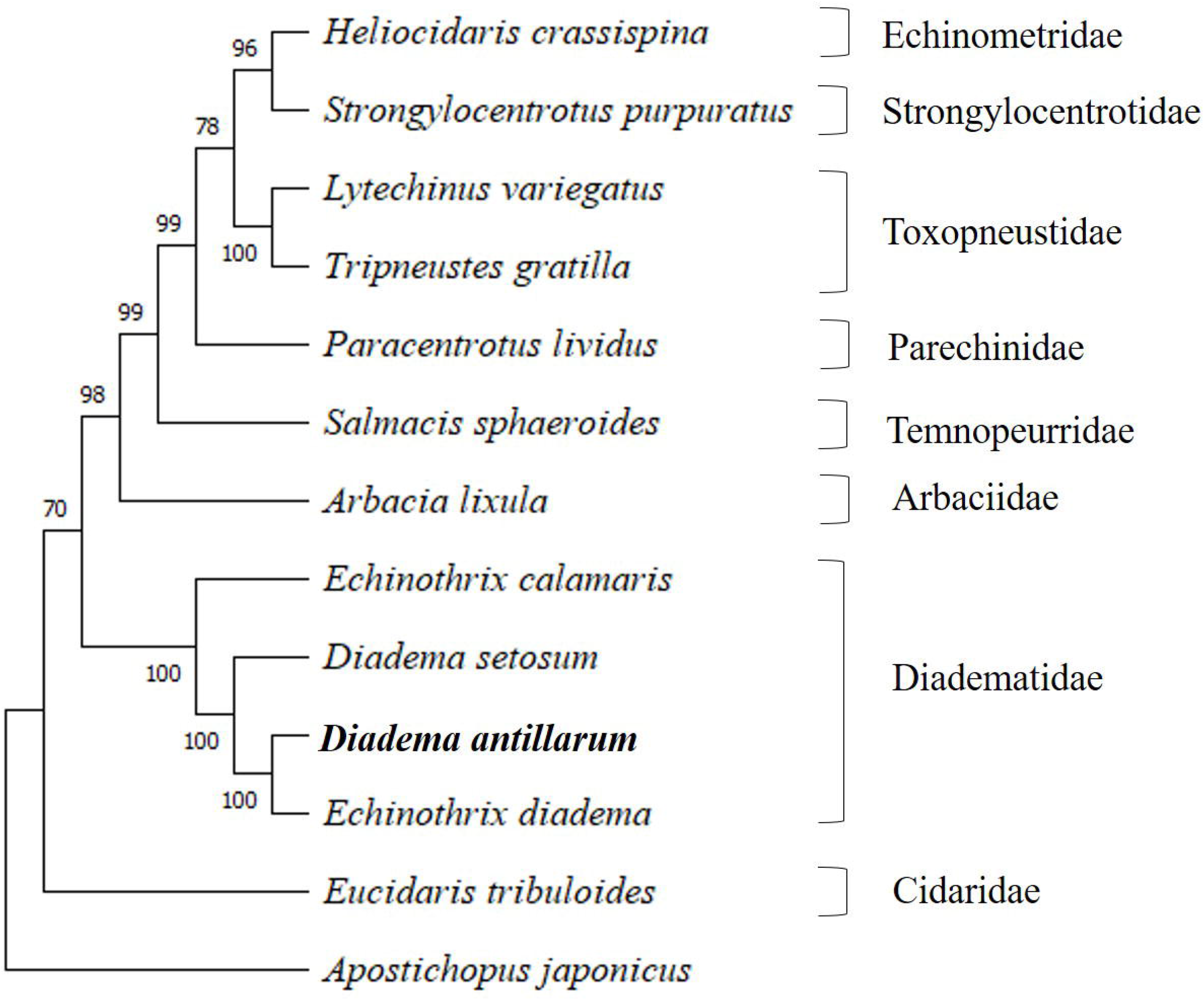
Phylogenetic analysis of mitochondrial genomes indicates that *Diadema antillarum* is more closely related to *Echinothrix diadema* than to *D. setosum*. A maximum likelihood consensus tree of sequences representing 12 sea urchin species and a sea cucumber (outgroup). Genbank accession numbers for taxa on the tree include: *H. crassispina* (KC479025.1, Jung, G. and Lee, Y.H., 2013, unpublished, direct submission), *S. purpuratus* (NC_001453.1, Valverde et al., 1996, Qureshi, S.A., and Jacobs, H.T. 1993, Jacobs et al., 1988), *L. variegatus* (NC_037785.1, Bronstein and Kroh 2018), *T. gratilla* (KY268294.1, Laruson 2017), *P. lividus* (J04815.1, Cantatore et al. 1989), *S. sphaeroides* (KU302103.1, Jung, G. and Lee, Y.H., 2013, unpublished, direct submission), *A. lixula* (X80396.1, De Giorgi et al., 1996), *E. calamaris* (NC_050274.1, Wakayama et al., 2019), *D. setosum* (KX385835.1, Li et al., 2016), *E. diadema* (KX385836.1, Li et al., 2016, unpublished direct submission), *E. tribuloides* (MH614962.1, Kroh A., and Bronstein, O., 2018, unpublished direct submission) and *A. japonicus* (NC_012616, Sun et al, 2010).

Interestingly, the tree topology shown in this study indicated that *D. antillarum* was more closely related to *E. diadema* than to *D. setosum*. The next most closely related taxon to these three was *E. calamaris*. For *E. diadema, E. calamaris* and *D. setosum*, the length of these mitochondrial genomes are 15,712 bp, 15,716 bp and 15,708 bp, respectively. A previous study reporting the complete mitogenome of *D. setosum* included a maximum likelihood phylogenetic tree with a highly supported sister clade (bootstrap value of 100) containing two taxa: *E. diadema* and *D. setosum* (Li et al., 2016). Furthermore, the results of our sequence identity matrix of mitogenomes between *D. antillarum* and *E. diadema* was 96.7% identical, whereas, *D. antillarum* vs. *D. setosum* or *D. antillarum* vs. E. *calamaris* was 86.3% and 80% identity, respectively. Additionally, the mitogenomes between *E. diadema* and *D. setosum* was 86.2% identical in sequence. One may suggest that the overall higher similarity between *D. antillarum* and *E. diadema*, as opposed to *D. setosum* may be due to the higher A + T bias in the mitogenome. However, the mitogenome sequences of *E. diadema* and *D. setosum* were generated from specimens collected from the South China Sea from the same research group (Li, Wu, Fu, and Zeng, 2016 unpublished, Li et. al., 2016). Given that there are other *Diadema* species in the South China Sea that are generally similar morphologically to *E. diadema* (including *D. setosum* and *D. savignyi*) the reported mitogenome of *E. diadema* may have actually been sampled from a *Diadema* species, namely *D. savignyi*. Unfortunately, the mitogenome has not been completed or otherwise available for *D. savignyi*. Nonetheless, to provide evidence for this claim we performed sequence analysis for three additional genes (*16S, ATPase* and *CO1*) for available data from species of the genera *Echinothrix* and *Diadema*. Results of the tree topologies for all three genes indicate that the specific gene sequences extracted from the mitogenome of *E. diadema* were more closely related to the particular gene sequences of *Diadema* spp. as opposed to those from *Echinothrix* spp. (**Supplemental Figures 2, 3, and 4**). In particular, the specific gene sequences that were extracted from the mitogenome of *E. diadema* were placed as a sister clade with *D. savignyi* (see Supplemental Figures 2, 3, and 4), which was placed next to a larger group of clades that included *D. africanum* (see Supplemental Figure 2), *D. antillarum* (see Supplemental Figures 2, 3, and 4) *D. mexicanum* (see Supplemental Figure 2) and *D. setosum* (see Supplemental Figures 2, 3, and 4). This was more distantly related to groupings of clades that included *E. diadema* (see Supplemental Figures 2) and *E. calamaris* (see Supplemental Figures 2, 3, and 4).

Furthermore, it is important to note that the habitat ranges of both *E. diadema* and *D. setosum* are in the Indo-Pacific region which is distinct from the habitat range of *D. antillarum*, which are typically found in the western Atlantic Ocean, the Caribbean Sea, the tropical coasts of South America down to Brazil, and from Bermuda to Florida. The authors are aware that a new subspecies of *D. antillarum*, called *D. antillarum ascensionis* has been recently identified in the eastern Atlantic, which is similar to the mid-Atlantic species of *D. antillarum* (Rodriguez, et al. 2013). This new subspecies is also similar to *D. africanum* that are found in the Eastern Atlantic islands, from Madeira Islands to the Guinean Gulf including Salvage Islands, Canary Islands, Cape Verde Islands (Hernández et al. 2008), and Sâo Tome Island (Lessios et al. 2001). While *D. antillarum* have been repopulating areas, which have extended their previous habitat range, we are not aware of any *D. africanum* that has migrated as far west as Puerto Rico, in the Caribbean Sea. In addition, when identifying the specimen prior to collection for this study, it lacked the typical iridophores that are identifiable in the sunlight of *D. africanum*. While the *ATPase* sequence analysis presented in this study does reflect that there are similarities in these sequence between *D. antillarum, D. africanum and D. mexicanum*, the *ATPase* gene sequence that was extracted from the mitogenome used in this analysis most closely resembles that of another *D. antillarum* (see Supplemental Figure 2). Thus, the authors think that the species used in this study has been correctly identified. Yet, as additional mitogenome sequences become available, the phylogenetic tree describing the relationships between genera within Diadematidae should become more thorough.

In conclusion, these results may provide some insight into the dispersal and speciation events of the family Diadematidae. However, more data is needed from other genera to draw further conclusions about the evolution and adaptation of this family lineage. The mitogenome sequence produced through this study can serve as a reference sequence for this species of sea urchin. In addition, the nuclear sequences generated in this study will be included in a larger study to assembly the whole genome for this species. Moreover, given the necessity of *D. antillarum* for maintaining the health and current structure of the remaining coral dominated coastlines in the face of global climate change, the genetics of this species will be likely important to include in future conservation and population genetics studies.

## 3. Data accessibility

The mitochondrial genome sequence has been deposited at DDBJ/ENA/GenBank under the accession number ON725136 (pending; submitted on May 19, 2022). The data is linked to the NCBI BioProject and BioSample numbers PRJNA839760 and SAMN28553754, respectively. All intermediate files are available in our GitHub repository: https://github.com/aglabx/mtDNA_assembly/tree/master/Diadema_anthilarum.

## Acknowledgements

Funding for the project was provided by start-up funds granted to Tarás Oleksyk by Oakland University.

