## Supplemental Figure 1 Legend for "The first complete mitochondrial genome of *Diadema antillarum* (Diadematoida, Diadematidae)"

**Supplemental Figure 1. Mitogenome assembly coverage map of *Diadema antillarum*.** A per-base coverage data table was generated using the Unipro UGENE software program. This graph was generated in Microsoft Excel (2019). The X-axis shows the base position of the mitogenome sequence. The Y-axis shows the assembly coverage at that base. The length of the mitogenome is 15,708 bps.
