## Supplementary figures and images for "The first complete mitochondrial genome of *Diadema antillarum* (Diadematoida, Diadematidae)"

### Supplemental Figure 1

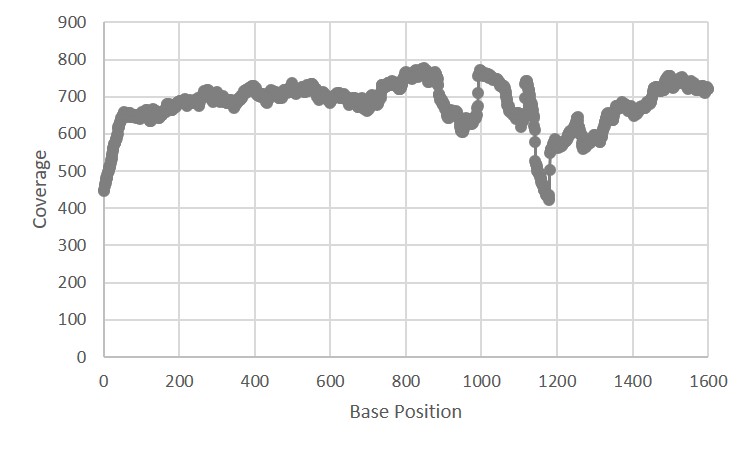

### Supplemental Figure 2

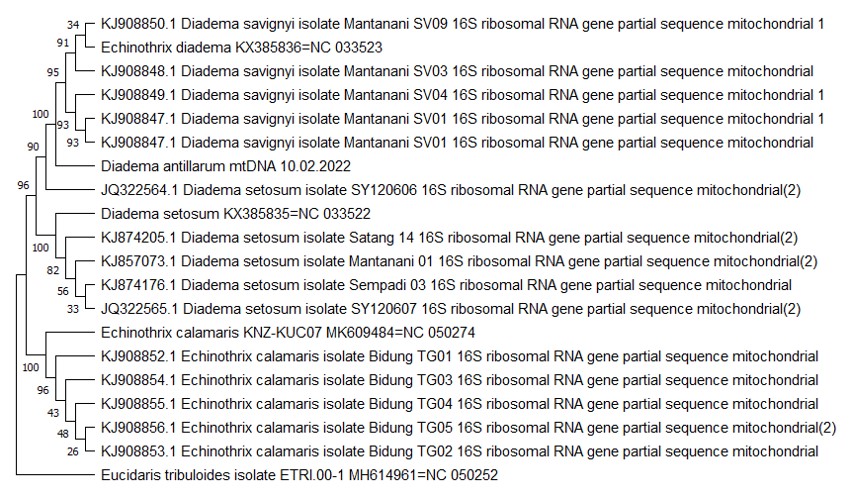

### Supplemental Figure 3

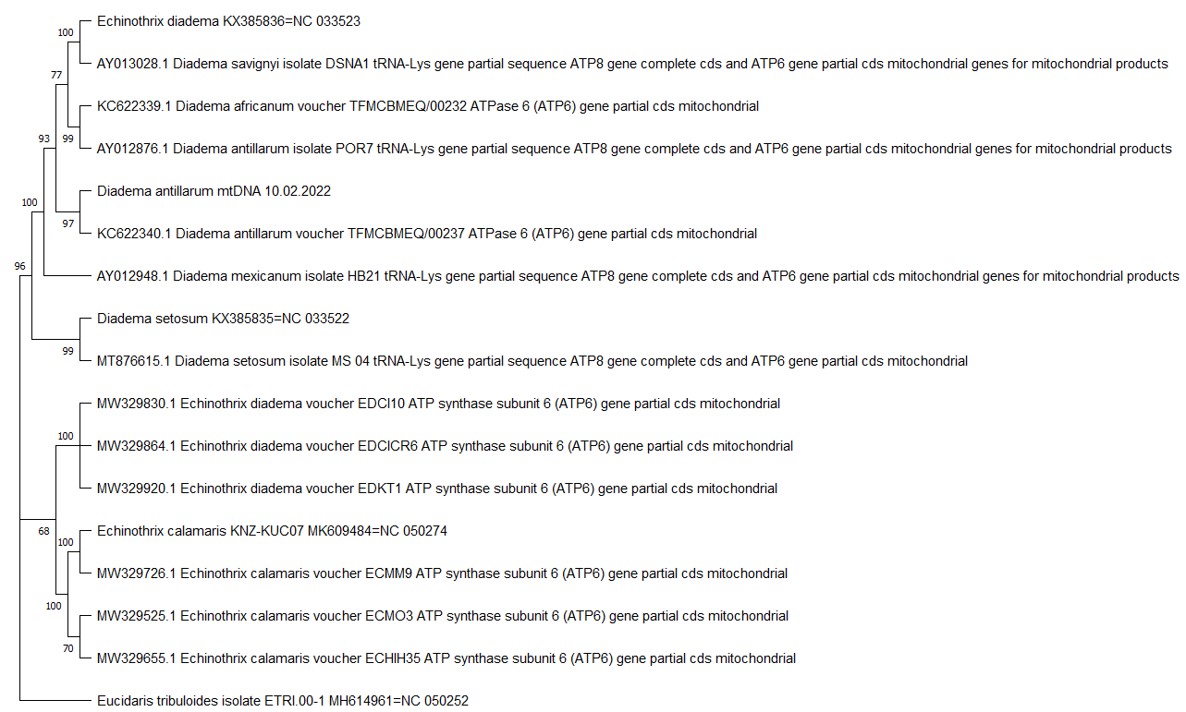

### Supplemental Figure 4

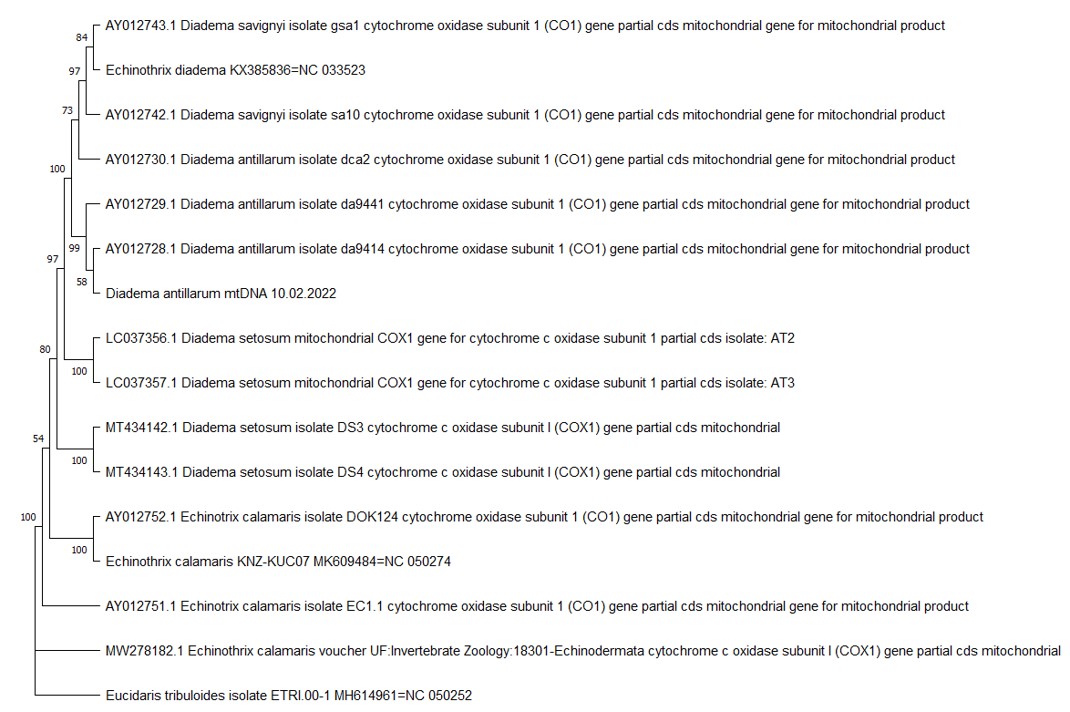
