## Supplemental Figure 2 Legend for "The first complete mitochondrial genome of *Diadema antillarum* (Diadematoida, Diadematidae)"

**Supplemental Figure 2. Phylogenetic analysis of 16S ribosomal RNA sequences for *Diadema and Echinothrix* spp**. A maximum likelihood bootstrapped consensus tree of partial gene sequences for 19 sequences of *Diadema and Echinothrix* spp. plus *Eucidaris tribuloides* as an outgroup. A section of the mitogenome for *E. diadema* (KX385836), *D. setosum* (KX385835) and *D. antillarum* (10.02.2022) that corresponded to the 16S gene was extracted by generating an initial alignment. To do this, the *E. calamaris* gene specific sequences listed in the tree were aligned to the entire *E. diadema* mitogenome. Then, the *D. setosum and D. savignyi* gene specific sequences listed in the tree were aligned to the entire *D. antillarum* mitogenome. Lastly, the *D. setosum* gene specific sequences listed in the tree were aligned to the entire *D. setosum* mitogenome. The rest of the mitogenome for each species was trimmed to only reflect the portion that pertained to the CO1 gene sequence. Thus, the taxa on the tree named Diadema antillarum mtDNA 10.02.2022, Echinothrix diadema KX385836=NC 033523, and Diadema setosum KX38535=NC 033522 include only the 16S gene sequences.
