## Supplemental Figure 3 Legend for "The first complete mitochondrial genome of *Diadema antillarum* (Diadematoida, Diadematidae)"

**Supplemental Figure 2. Phylogenetic analysis of ATPase genes** **for *Diadema and Echinothrix* spp**. ATP synthase subunit 6 gene (partial coding sequence) and ATP synthase subunit 8 gene (complete coding sequence) were included in the analysis, as well as tRNA-Lys gene (partial sequence), which was included in some of the taxa. A maximum likelihood bootstrapped consensus tree is shown for 16 sequences of *Diadema and Echinothrix* spp. plus *Eucidaris tribuloides* as an outgroup. A section of the mitogenome for *E. diadema* (KX385836), *D. setosum* (KX385835), *E. calamaris* (KNZ-KUC07) and *D. antillarum* (10.02.2022) that corresponded to the specific gene was extracted by generating an initial alignment. To do this, the *E. calamaris* gene specific sequences listed in the tree were aligned to the entire *E. diadema* and *E. calamaris* mitogenomes. Then, the *D. antillarum* gene specific sequences listed in the tree were aligned to the entire *D. antillarum* mitogenome. Lastly, the *D. setosum* gene specific sequences listed in the tree were aligned to the entire *D. setosum* mitogenome. The rest of the mitogenome for each species was trimmed to only reflect the portion that pertained to the ATPase gene sequences. Thus, the taxa on the tree named Diadema antillarum mtDNA 10.02.2022, Echinothrix diadema KX385836=NC 033523, Diadema setosum KX38535=NC 033522 and Echinothrix calamaris KNZ-KUC07 MK609484=NC050274 include only the ATPase gene sequences.
