## Supplemental Figure 4 Legend for "The first complete mitochondrial genome of *Diadema antillarum* (Diadematoida, Diadematidae)"

**Supplemental Figure 4. Phylogenetic analysis of cytochrome oxidase 1 (*CO1*) genes** **for *Diadema and Echinothrix* spp**. A maximum likelihood bootstrapped consensus tree is shown for 15 partial coding sequences of *Diadema and Echinothrix* spp. plus *Eucidaris tribuloides* as an outgroup. A section of the mitogenome for *E. diadema* (KX385836) and *D. antillarum* (10.02.2022) that corresponded to the CO1 gene was extracted by generating an initial alignment. To do this, the *E. calamaris* gene specific sequences listed in the tree were aligned to the entire *E. diadema* and *E. calamaris* mitogenome. Separately, the *D. antillarum* gene specific sequences listed in the tree were aligned to the entire *D. antillarum* mitogenome. The rest of the mitogenome for each species was trimmed to only reflect the portion that pertained to the CO1 gene sequence. Thus, the taxa on the tree named Diadema antillarum mtDNA 10.02.2022 Echinothrix calamaris KNZ-KUC07 MK609484=NC050274 and Echinothrix diadema KX385836=NC 033523 include only the CO1 gene sequences.
